# Notes on the endemic cloud forest plants of the Cameroon Highlands and the new, Endangered, *Tricalysia elmar* (Coffeeae-Rubiaceae)

**DOI:** 10.1101/709303

**Authors:** Martin Cheek, Isla Causon, Barthelemy Tchiengue, Eden House

## Abstract

**Background and aims:** This paper reports a further discovery in the context of a long-term botanical survey in the Cross River-Sanaga interval of west-central Africa, focussing on species discovery and conservation.

**Methods:** Normal practices of herbarium taxonomy have been applied to study the material collected. The relevant collections are stored in the Herbarium of the Royal Botanic Gardens, Kew, London and at the Institute of Research in Agronomic Development – National Herbarium of Cameroon.

**Key results:** New species to science continue to be discovered from the cloud (submontane) forests of the Cameroon Highlands in Cameroon. Most of these species are rare, highly localised, and threatened by habitat destruction. These discoveries increase the justification for improved conservation management of surviving habitat. *Tricalysia elmar* Cheek (*Coffeeae*-Rubiaceae) is described as an additional species new to science and is compared with similar species of the genus. Restricted so far to four locations, Mt Kupe, Bali Ngemba, Lebialem and Nta Ali, its conservation status is assessed as Endangered (EN B1+2ab(iii)) according to the 2012 criteria of IUCN.

## INTRODUCTION

The new species of *Tricalysia* reported in this paper was discovered as a result of the long-term survey of plants in Cameroon to support improved conservation management led by botanists from Royal Botanic Gardens, Kew and the IRAD (Institute for Research in Agronomic Development)-National Herbarium of Cameroon, Yaoundé. This study has focussed on the Cross-Sanaga interval (Cheek et al. 2001) which contains the area with the highest species diversity per degree square in tropical Africa (Barthlott et al. 1996), including several endemic genera. *Medusandraceae* sensu stricto (*Medusandra* Brenan) was until recently considered endemic to the Cross-Sanaga interval (Heywood et al. 2007) but data now shows that *Medusandra*, with *Soyauxia* Oliv., are confamilial with neotropical *Peridiscaceae* (Soltis et al. 2007, Breteler et al. 2015).

The herbarium specimens collected in these surveys formed the primary data for the series of Conservation Checklists that began at Mt Cameroon (Cheek et al. 1996), with the Plants of Mt Cameroon (Cable & Cheek 1998) and continued with Mt Oku and the Ijim Ridge (Cheek et al. 2000), Bali-Ngemba (Harvey et al. 2004), the Mt Kupe, the Bakossi Mts and Mwanenguba (Cheek et al. 2004), Dom (Cheek et al. 2010) and Lebialem (Harvey et al. 2010). So far, over 100 new species and several new genera have been discovered and published as a result of these surveys, many from the cloud forest altitudinal band of 800– 2000 m altitude, and two new national parks have resulted. The species described in this paper was first noted as “a new species, apparently endemic to Mt Kupe and Bali Ngemba” in Cheek et al. (2004: 394) and in Harvey et al. (2004: 124), and was subsequently found at Lebialem (Harvey et al. 2010: 142) and was given the provisional name of “*Tricalysia sp. B aff. ferorum”.* Here it is formally named as *Tricalysia elmar* Cheek, allowing IUCN to accept a conservation assessment for the species, and for it then to be incorporated into conservation prioritisation initiatives.

### The genus *Tricalysia* A.Rich

The new species was readily placed in *Tricalysia* of tribe *Coffeeae* by the combination of shortly sheathing and cylindrical interpetiolar stipules with long awns, contracted axillary inflorescences with a series of 3(–4) cupular calyculi subtending the flowers, calyx teeth and tube conspicuous, corolla tubes about as long as lobes, anthers largely exserted, inserted at the mouth of the corolla tube which lacks exserted hairs.

The genus *Tricalysia* currently contains about 78 described species, which are mainly evergreen hermaphroditic shrubs or small trees of lowland evergreen forest in tropical and southern Africa and Madagascar. However, several species are subshrubs of drier habitats in southern Africa, and at least one species is deciduous. Formerly, the name *Tricalysia* was extended to Asian species now included in *Diplospora* DC. and *Discospermum* Dalzell (Ali & Robbrecht 1991). Those Asian species are distinct from *Tricalysia* s.s. (in Africa) by their unisexual tetramerous flowers. *Tricalysia* is now restricted to Africa and Madagascar. *Sericanthe* Robbr., an African genus segregated from *Tricalysia* by Robbrecht (1978), is distinguished by possessing bacterial nodules in the leaves and a sericeous indumentum on the outer corolla surface. The molecular phylogenetic analysis of *Tricalysia* by Tosh et al. (2009) showed that the former *Tricalysia* subgenus *Empogona* had a sister relationship with *Diplospora*. Accordingly, the African genus *Empogona* Hook.f. was resurrected. Its species are distinguished from *Tricalysia* by black fruits, flag-like anther connectives and free, often alternate distal bracts. Most African *Tricalysia* species are well studied as a result of Robbrecht’s revisions (Robbrecht 1979, 1982,1983,1987). Within *Tricalysia* as currently defined, Robbrecht (1979, 1982,1983,1987) recognised four sections in Africa: Sect. *Ephedrantha* Robbr., Sect. *Probletostemon* (K.Schum.) Robbr., Sect. *Tricalysia* and Sect. *Rosea* (Klotzsch) Robbr. However, only eleven species of the 78 accepted *Tricalysia* were sampled in the phylogenetic study by Tosh et al. (2009), and of these, only six were African, so further, more intensively sampled, phylogenetic studies are called for if the current sectional classification is to be tested. Since the monographic studies of Robbrecht, seven new taxa have been published from the Flora Zambesiaca region by Bridson (in Bridson & Verdcourt 2003) and nine new taxa published for Madagascar (Sect. *Androgyne* Robbr.) by Ranarivelo-Randriambovanjy et al. (2007). Apart from these, only two other new species have been published, both from Cameroon: *Tricalysia lejolyana* Sonké & Cheek (Sonké et al. 2002a) and *T. achoundongiana* Robbr., Sonké, & Kenfack (Sonké et al. 2002b). However, new species are likely to be discovered in Gabon, where 118 *Tricalysia* specimens are recorded as being unidentified to species versus 219 identified (Sosef et al. 2005: 373–375), and in Cameroon, where six unidentified, possibly undescribed species are recorded from the Bakossi area in Cheek et al. (2004). Many species of the genus are geographically localised, rare and threatened. For example, twelve species of *Tricalysia* were assessed as threatened in Cameroon (Onana & Cheek 2011), mainly because they are known from a single or few locations, and some are threatened by logging followed by agriculture such as palm oil plantations, e.g. *Tricalysia lejolyana* (Sonké et al. 2002a).

*Tricalysia elmar* is distinguished by having fruit with a highly conspicuous, large and dome-like accrescent disc. This feature is otherwise only known in the genus in *T. ferorum* Robbr., placed in Sect. *Probletostemon* due to its free distal bracts, yet placed there with reservations: “of isolated position, does not fully match with the enumerated characters, but is included here” (Robbrecht 1983: 299). However, *Tricalysia elmar* has distal bracts united in a cupular calyx, and due to this character, together with the entire calyx, stalked and entirely exserted stamens, it merits placement in Sect. *Tricalysia*. In the key to the species of this section (Robbrecht 1987: 67–71), *Tricalysia elmar* keys out to couplet 44, due to the calyx lobes exceeding the tube in length, the 6-merous flowers, glabrous styles and anthers, pubescent ovary and corolla lobes 8–11 mm long. However its characters fit neither of the two choices offered in that couplet, which lead to *T. niamniamensis* Hiern and, in couplet 45, to *T. bagshawei* S.Moore and *T.oligoneura* K.Schum. *Tricalysia elmar* differs from the first species in lacking petal lobes with entirely pubescent abaxial surfaces (instead they are only sparsely hairy at the base and apex of the lobes). It differs from the last two species in lacking hairs at the base of the corolla tube (they are present only at the apex), and lacking pubescence only on half of the corolla lobes not covered in bud (hairs present are spread equally on both sides of the lobe). Below we present a table using diagnostic characters separating *Tricalysia elmar* from these species, drawing on data from Robbrecht (1983, 1987).

A key to the 12 genera in tribe *Coffeeae* can be found in Cheek et al. (2018a). This tribe has its highest generic diversity in Cameroon with nine of the genera present.

## MATERIAL AND METHODS

The methodology for the surveys in which this species was discovered is recorded in Cheek & Cable (1997). Nomenclatural changes were made according to the Code (Turland et al. 2018). Names of species and authors follow IPNI (continuously updated). Herbarium material was examined with a Leica Wild M8 dissecting binocular microscope fitted with an eyepiece graticule measuring in units of 0.025 mm at maximum magnification. The drawing was made with the same equipment with a Leica 308700 camera lucida attachment. Specimens were inspected from the following herbaria: BM, K, P, WAG, YA. The format of the description follows those in other papers describing new species in *Coffeeae* e.g. Cheek et al. (2018a). Terminology for specialised structures, e.g. for colleters, and domatia, generally follows Robbrecht (1987, 1988). All specimens cited have been seen unless indicated “n.v.”. The conservation assessment follows the IUCN (2012) categories and criteria. GeoCAT was used to calculate red list metrics (Bachman et al. 2011). Herbarium codes follow Index Herbariorum (Thiers, continuously updated).

## RESULTS

The new species, *Tricalysia elmar*, shares uniquely within the genus an accrescent floral disc with *T. ferorum* of Sect. *Probletostemon.* The disc in both species conspicuously occupies the apex of the fruit. In other species of the genus the disc is not accrescent and is usually invisible in fruit, where it is concealed by the calyx. However, the taxonomic placement of the new species is postulated as being not with Sect. *Probletostemon* but with species of Sect. *Tricalysia*, since it has the following traits, that characterise that group (Robbrecht 1987) yet which are unknown in Sect. *Probletostemon* (Robbrecht 1983).

1. distal bracts united in a cupular calyculus;
2. calyx entire, not spathaceous or split;
3. anthers completely exserted from the corolla mouth, with distinct filaments;
4. hairs of the corolla tube not exserted from the corolla mouth.

Using the dichotomous key of Robbrecht (1987), *Tricalysia elmar* keys out to the vicinity of three species (see introduction). Its affinities are therefore likely to be with these, although molecular phylogenetic studies are needed to confirm this hypothesis. The taxa can be separated using the characters shown in table 1.

**Table 1.** Characters separating *Tricalysia elmar* from similar species. Data for *Tricalysia ferorum* from Robbrecht (1983) and for *T. niamniamensis*, *T. bagshawei*, and *T. oligoneura* from Robbrecht (1987).

|  | *Tricalysia*<br>*ferorum* | *Tricalysia*<br>*elmar* | *Tricalysia*<br>*niamniamensis* | *Tricalysia*<br>*bagshawei* | *Tricalysia*<br>*oligoneura* |
| --- | --- | --- | --- | --- | --- |
| **Distal bracts**<br>**free or united**<br>**in calyculus**<br>**cup** | Free | United in cup | United in cup | United in cup | United in cup |
| **Corolla tube**<br>**inside densely**<br>**hairy at least in**<br>**one band near**<br>**stamen**<br>**insertion** | Densely hairy | Dense hair<br>band | Glabrous or with<br>few hairs | Densely hairy | Densely hairy |
| **Indumentum**<br>**on adaxial**<br>**surface of**<br>**corolla lobes** | Glabrous<br>apart from<br>ciliate margin | Sparse hairs<br>only at base in<br>apex | Densely<br>pubescent<br>throughout | Pubescent only<br>on half exposed<br>in bud | Pubescent only<br>on half exposed<br>in bud |
| **Disc**<br>**conspicuous,**<br>**dome-like,**<br>**occupying**<br>**much of apex**<br>**of fruit** | Conspicuous | Conspicuous | Inconspicuous | Inconspicuous | Inconspicuous |
| **Petiole length**<br>**(mm)** | 2.5 | 5–12 | 1–6 | (6–)10–15 | 1–3 |
| **Leaf-blade**<br>**shape** | Obovate | Elliptic-oblong (-<br>oblanceolate<br>or lanceolate) | Ovate, elliptic or<br>obovate | Ovate, elliptic<br>or obovate | Ovate, elliptic<br>or obovate |
| **Leaf-blade**<br>**dimensions**<br>**(cm)** | (4–)6–11(–<br>13) x (2.5–<br>)3.5–5(–7.5) | 12–19.8(–<br>21.2) x 4.5–<br>7.4 | (2.5–)4.5–9(–12)<br>x (1–)1.5–2.5(–3) | (3–)5–9(–11) x<br>(1.5–)2.4(–4.7) | 8–17(–20) x<br>(2.5–)6.5(–8) |
| **Secondary**<br>**nerve number**<br>**on each side of**<br>**midrib** | 3–5(–6) | 6–10 | 4–6 | 3–4(–5) | 3–5(–6) |
| **Distribution** | Cameroon | Cameroon | DRC, Kenya,<br>Uganda, to<br>Zimbabwe | Uganda,<br>Tanzania, DRC,<br>Zambia | Nigeria to DRC,<br>CAR |
| **Altitude range** | <800 m | 1310–1850 m | To 1530 m | To 1200 m | To 800 m |

### *Tricalysia elmar* Cheek, sp. nov

Similar to *Tricalysia ferorum* Robbr. of Sect. *Probletostemon* (K.Schum.) Robbr. in the large, conspicuous, dome-like accrescent disc in the fruit, differing in the entire, united distal calyculus (distal bracts not free) and the glabrous style and anthers (not hairy); differing from all species of Sect. *Tricalysia* in the dome-like accrescent disc in the fruit. – Type: Cameroon, South West Region, Mount Kupe Summit, Max’s trail, 4°49’N 9°43’E, 1550m alt., fl. 2 Nov. 1995, Cheek 7619 (holo-: K; iso-: BR, MO, P, SCA, YA).

*Tricalysia sp. B. aff. ferorum* Cheek in Cheek *et al.* 2004: 394; Harvey *et al.* 2004: 124; Harvey *et al.* 2010: 142.

<u>Evergreen tree</u>, rarely a shrub, (3–)7–13 m tall, trunk brown; flowering and fruiting stems at base (proximal to trunk), matt whitish brown, lacking lenticels, not or sparingly branched, 7–8 mm diam., glabrous; third internode from stem apex (2–)3–5(–6) mm diam.; indumentum at stem apex dense (covering c.95% of surface) with yellow-green appressed hairs 0.1–0.4 mm long, cover declining to 50–70% cover at the third internode from the apex, apical bud sometimes with an amber coloured bead of exudate; internodes (1.5–)4–5(–7) cm long, (3–)5– 6 pairs of leaves present per stem, leaves subequal in shape and size. <u>Stipules</u> persistent, 1–5 x 4–7 mm, shortly sheathing, limb broadly cylindric, distal part broadly triangular, awns apical, 3–6 x 1–2 mm, laterally compressed, outer surface indumentum as stem apex, inner surface c. 95%. covered in hairs c. 0.5 mm long; colleters reduced standard type, lacking feathery appendages, narrowly cylindrical 0.3–0.5 x 0.03 mm, amber coloured, inserted in line at inner base of stipule, inconspicuous amongst the dense indumentum. <u>Leaf-blades</u> elliptic-oblong, less usually oblanceolate or lanceolate, 12–19.8(–21.2) x 4.5–7.4 cm, apex acuminate, acumen 0.6–1.5(–1.8) x (0.1–)0.4–0.8 cm, base acute to cuneate, slightly decurrent down the petiole, leaf margin brochidodromous, upper surface drying dark grey-brown, lower surface pale brown, secondary nerves 6–10 on each side of the midrib, midrib and secondary nerves deeply impressed on upper surface, domatia tuft type, shallowly excavated, sparsely hairy (Fig. 1B), hairs 0.15–0.2 mm long. Petiole shallowly canaliculate, 0.5–1.2 x 0.1–0.2 cm, indumentum as stem apex. <u>Inflorescences</u> single, in both axils of a node, fertile nodes per stem 1–3, flowers per inflorescence 4(–7), calyculi 3(–4), cup-shaped, more or less completely concealing the axis, indumentum as stem apex; 1^st^ order (proximal) calyculus 1.1– 1.3 x 2.8–3.5 mm, divided half way to base on each side, rarely with foliar lobes as 2^nd^ order calyculi, subtending 1–2 lateral flowers, supplementary calyculus subtending additional flower (s) sometimes present; 2^nd^ order (medial) calyculus, (1.1–)1.8–2 x 2.7–3.5 mm, foliar lobes two, spatulate or lanceolate, 1.1–5.1 x 0.4–1.6 mm, stipular lobes if present, 1–2, triangular, 0.5–1.0 x 0.3–1.3 mm, subtending 3 flowers, each with a single 3^rd^ order (distal) calyculus; 3^rd^ order calyculi, 1.6–2.2 x (2.2–)2.8–3.1 mm, entire (Fig. 1F) or lobed, lobes 4–6, shallow, 0.4–0.5(–1.2) x 1.1–1.3(–1.6) mm; pedicels absent or minute, <0.1 mm long, concealed by calyculus. <u>Flowers</u> hermaphrodite, homostylous, 6-merous, sweetly scented (*Etuge* 4690). Ovary-hypanthium 1–3 x 2–4 mm, base concealed within 3^rd^ order calyculus, calyx tube 0.6–1.3 mm long; calyx lobes triangular, stout, erect, 1–2 x 1 mm, outer indumentum as stem apex but hairs 0.1–0.2 mm long, inner surface less densely hairy, colleters not seen. Corolla white, tube 3–6 x 2–3 mm, dilating to 5 mm wide at mouth, glabrous outside, inside with hair band 2–5 mm wide, inserted below mouth of tube, hairs 0.3–0.4 mm long, appressed, otherwise glabrous. Petal lobes oblong, (8–)10–11 x 1–3 mm, petal apex acute, often laterally compressed (Fig. 1E) sparsely hairy, hairs 0.04–0.1mm long, erect, on margin and base and apex of abaxial surface. Stamens usually fully exserted, 6–7.5 mm long, anthers oblong, 5–8 x 0.6–1.2 mm, submedifixed, apical connective appendage 0.2– 0.4 x 0.1–0.2(–0.4) mm, usually hooked; filaments 0.5–1.5 mm long, inserted below mouth of tube at 1.3–1.5 mm, triangular, c. 0.2 mm wide at base, tapering to 0.1 mm wide at apex, glabrous. Disc subcylindrical, apex concave, rarely flat, 0.2–0.5 x 0.8–1.4 mm, glabrous. Style 9.5–9.6(–14) x 0.3–0.5 mm, exserted, glabrous, bifurcating into two linear stigmatic lobes, at length diverging, stigmatic lobes (1.4–)1.9 mm long, apex tapering, glabrous. Ovary 2–locular, ovules 3 per locule. <u>Fruit</u> dark green and hairy when immature, orange or yellow and sparsely hairy at maturity (*Tchiengue* 1915), fruit wall 0.8–1.4 mm thick, globose or shortly ellipsoid (Fig. 1I), (9–)11–13(–17) x 8–13(–16) mm, crowned by a persistent calyx 4– 10 mm diam., calyx lobes 1–2 x 1–3 mm, inner surface of calyx densely hairy (Fig. 1J); infructescence axis not accrescent, 3–7 mm long (Fig. 1I). Disc persistent, strongly accrescent, conspicuous, dome-like, 4–7 mm diam., projecting 2 mm from the calyx, glabrous. <u>Seeds</u> (3–)4–5 per fruit, elliptic in profile, laterally compressed, 7–8 x 3–4 x 2–3 mm, seed coat glossy, hard, dark brown, and undulate surface, hilum matt, pale brown, extending the length of the seed along one edge, 0.5–0.75 mm wide. Fig. 1 & 2.

**Figure 1.**
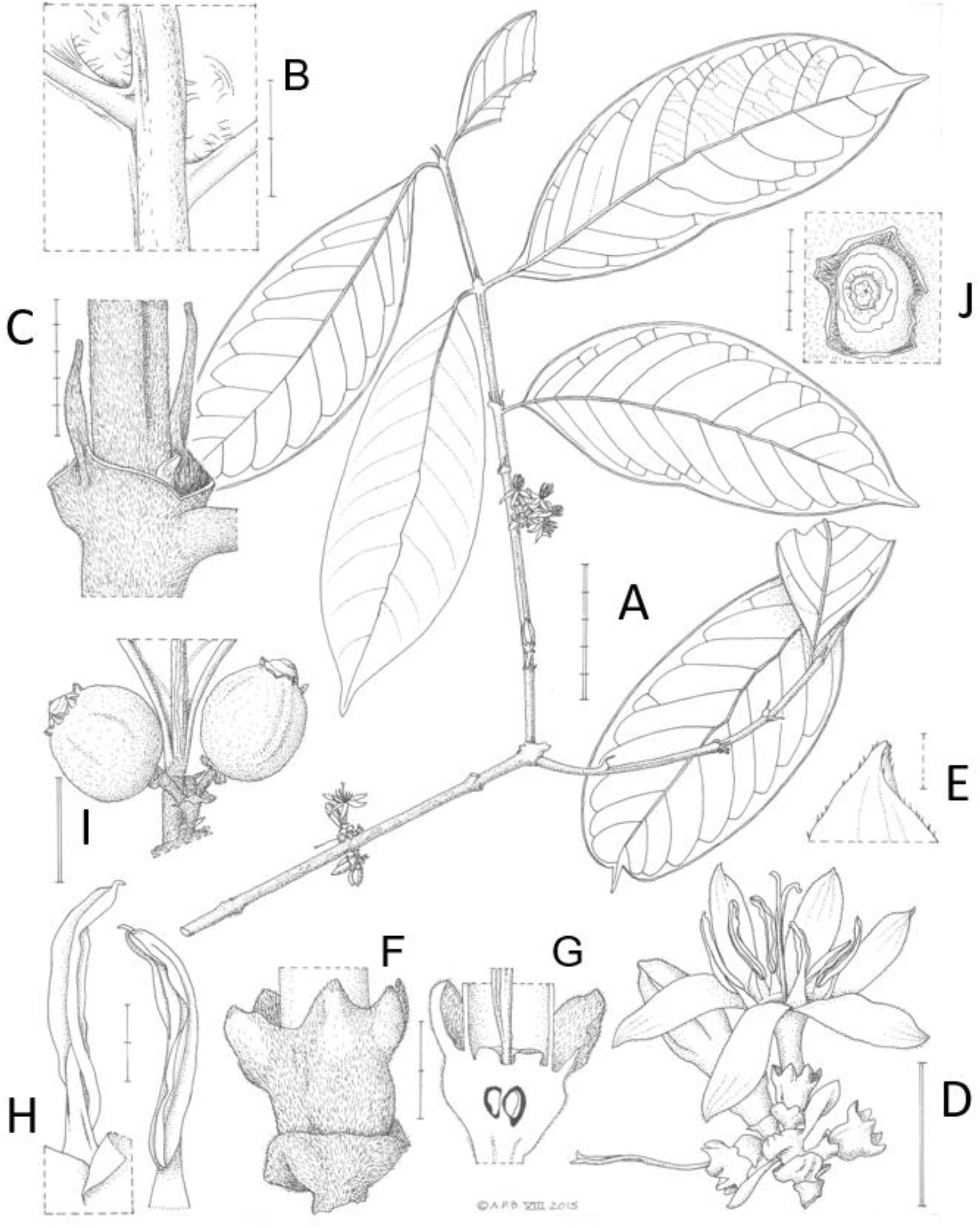
*Tricalysia elmar*: A, habit; B, domatia on leaf abaxial surface; C, stipules; D, inflorescence with calyculi, foliar and stipular lobes, flower bud, open flower and flower after corolla drop; E, petal tip showing scattered hairs; F, calyx and 3^rd^ order (distal) calyculus; G, longitudinal section of flower showing disc, ovary, and base of style; H, stamens, outer face with anther lobe bases (left) and inner face (right); I, fruits in situ; J, fruit apex showing calyx surrounding accrescent disc. Scale bars: A = 5 cm; B, H, F, G = 2 mm; C, J = 5 mm; D, I = 1 cm; E = 500 µm. A-H from *Cheek* 7619 (K); I and J from *Etuge* 1735 (K), all drawn by Andrew Brown.

**Figure 2.**
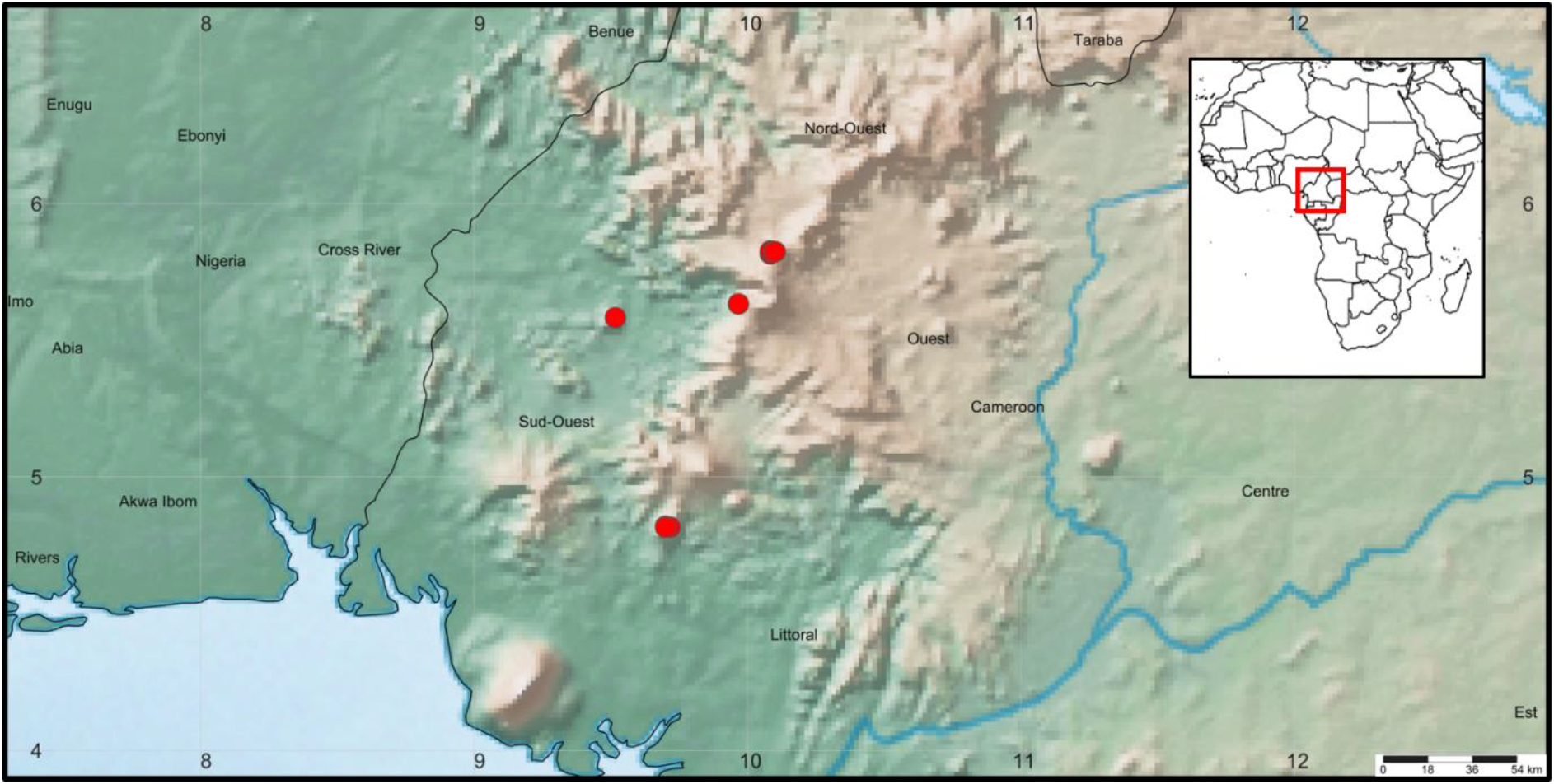
*Tricalysia elmar* global distribution (marked in red).

### Habitat and distribution

Cameroon, known only from Mt Kupe, Mt Nta Ali, Lebialem Highlands of SW Region, and Bali Ngemba Forest reserve of NW region.

Tree of upper cloud (submontane) forest; 1310–1850 m alt.

### Available specimens examined

**Cameroon: South West Region. Mt Kupe:** Nyasoso, Max’s trail, 1600m alt., fr., 6 June 1996, *Etuge et al*. 2176 (BR, K, MO, P, SCA, WAG, YA); Mount Kupe Summit, Max’s trail, 4°49’N 9°43’E, 1550m alt., fl., 2 Nov. 1995, *Cheek* 7619 (K, MO, P, SCA, WAG, YA); Nyasoso, Max’s trail to Mt. Kupe, 1700m alt., fr., 27 Feb. 1996, *Etuge* 1735 (K, MO, SCA, WAG, YA); Kupe Village, path from Kupe village to summit, 1600m alt., fr., 23 May 1996, *Cable et al*. 2559 (B, K, MO, P, SCA, WAG, YA); Nyasoso, Earthwatch peak 1, 1850m alt., fr., 6 June 1996, *Cable et al*. 2926 (K, MO, P, SCA, WAG, YA). **Nta Ali Forest Reserve**: 5.°35.00’N 9.°31.00’E, c. 800–1250m alt., fr., 12 Mar. 1995, *Thomas* 10523 (K, YA, WAG, MO, P); **Lebialem Highlands:** Fosimondi, 5°38’00.0’N 9°58’00.0’E, 1800m alt., fr., 16 Apr. 2004, *Tchiengue et al*. 1915 (K, YA n.v.). **North West Region. Bali Ngemba Forest Reserve:** Mantum, 5°49’00.0’N 10°5’00.0’E, 1400m alt., fl., 8 Nov. 2000, *Etuge et al*. 4690 (K, YA n.v.); Bali Ngemba Forest Reserve, 5°49’35.0’N 10°5’00.0’E, 1500m alt., fl., 9 Nov. 2000, *Tadjouteu et al*. 399 (K, YA n.v.); Mantum, 5°49’32.0’N 10°5’42.0’E, 1310m alt., fr., 11 April 2002, *Zapfack et al*. 2016 (K, YA n.v.); Mantum, 5°49’27.0’N 10°6’04.0’E, 1700m alt., fr., 12 April 2002, *Zapfack et al*. 2046 (K, YA n.v.).

### Etymology

Named (noun in apposition) for Prof. Elmar Robbrecht (1946-) of the Meise Botanic Garden and Herbarium, Belgium, the noted global specialist in Rubiaceae who monographed and laid the systematic foundations for all future research on the genus *Tricalysia*.

### Conservation

We calculated with GeoCAT (Bachman et al. 2011) the extent of occurrence as 3,173 km^2^ and the area of occupancy as 96 km^2^ using the IUCN preferred 4 km^2^ grid cells. There are four locations, with major threats to the cloud forest habitat of *Tricalysia elmar* present at three of these. At Mt Kupe, clearance of forest upslope from Nyasoso for subsistence agriculture from the expanding human population continues unabated, and the designated protected area at the summit has not yet been formally recognised (Cheek et al. 2004, Etuge pers. comm. 2019). At Lebialem, clearance downslope from villages on the high plateau for agricultural land also continues apace (Harvey et al. 2010, Tchiengue pers. obs. 2008-present). At Bali Ngemba, timber extraction followed by forest clearance for agriculture both downwards from villages on the high plateau above, and upwards from the direction of Bali, also continue (Harvey et al. 2004, Tah pers. comm. 2019) and are steadily degrading the quality and area of this isolated natural forest fragment. The species only seems secure at the Nta Ali location, where there is currently no evidence of threats. Therefore, we assess *Tricalysia elmar* as Endangered, (EN B1+B2ab(iii)). The absence of the species from apparently suitable habitat at Mt Etinde, in the Bakossi Mts, Rumpi Hills and other sites in the Cameroon Highlands may be due to under-collection, but is just as likely to reflect reality of absence at sites such as Mt Etinde which are relatively well-surveyed. It is to be hoped that the locations for this species will become formally protected (at present none are) for conservation and supported by local communities, otherwise as things stand currently, this species is at risk of extinction at three of its four known locations

### Notes

At Bali Ngemba, *Tricalysia elmar* (as *T. sp. B aff. ferorum*) is one of the 14 species that characterise the vegetation type “submontane forest with *Pterygota mildbraedii* (1300– 1700 m)” (Harvey et al. 2004: 17).

At Mt Kupe it is one of the few tree species restricted to the upper submontane forest (1400– 1900 m altitude). The number of species restricted to lower submontane forest (800–1400 m attitude) or occurring throughout the submontane belt, is far higher (Cheek et al. 2004: 36– 37). This taxon is also listed as one of the 33 endemic and near-endemic taxa occurring at Mt Kupe (Cheek at al. 2004: 39).

### The Accrescent disc in Rubiaceae

The floral disc in Rubiaceae e.g. in *Tricalysia*, is considered to secrete nectar for the attraction of pollinators (Robbrecht 1987: 53). Therefore, it is not unexpected that as the ovary develops into the fruit, it is generally invisible or inconspicuous, concealed by the calyx, having no obvious function at that stage. However, several species of Rubiaceae in a range of genera and tribes are distinguished from their congeners at the fruiting stage by the presence of accrescent discs which are highly conspicuous to the naked eye, occupying much of the apex of the fruit. Elsewhere in the *Coffeeae* this feature can be seen in *Coffea bakossii* where it helps to distinguish it from the similar *C. liberica* Hiern (Cheek et al. 2002). In the Vanguerieae, massive accrescent discs are an important character for recognising *Keetia susu* Cheek and *Keetia abouabou* Cheek (Cheek et al. 2018d) from other West African species of the genus. In all these examples, those species with accrescent discs are distinguished by having larger fruits than their congeners suggesting that discs accrescence is linked to the development of larger fruit size which can be conjectured to have selective dispersal advantage in some scenarios, where animal dispersers might favour larger fruits.

### Cloud, or submontane forest in the Cameroon Highlands

A fault, running nearly NE-SW between two major African plates, is the origin of the Cameroon Highlands that lie in a band 50–100 km wide along that fault and which were formed over four separate periods of mountain building beginning in the Tertiary, although a Cretaceous origin has also been suggested. The highest point, at 4095 m, is Mt Cameroon, an active hawai`ian type volcano, but the highlands begin in the bight of Biafra 40 km to the SW with the mountain-island of Bioko (formerly Fernando Po), part of Equatorial Guinea. Bioko is geologically and botanically nearly a twin of Mt Cameroon. Moving inland along the line, are the Rumpi Hills to the northwest, with Nta Ali as their northern outlier. NE of Mt Cameroon and its subsidiary peak Mt Etinde (1700 m alt.), are Mt Kupe (2064 m alt.), the Bakossi Mts (1895 m), and further northeast the Mwanenguba caldera (2411 m). Further along the line are the Bamileke Plataeau, and the high lava plateau of the Bamenda Highlands (2000 m alt.), including Mt Oku (3011 m alt.) and the Kilum Ridge the second highest peak in Cameroon. The highlands branch northwestwards into Nigeria in two places, as the Obudu plateau, and from the Bamenda Highlands, the Mambilla Plateau. Continuing northeastwards to Tchabal Mbabo, the highlands turn eastwards as the Adamaoua Plateau, extending into the west of the Central African Republic (Courade 1974, Cable & Cheek 1998, Cheek et al. 2004). Moisture laden southwesterly monsoon winds from the Atlantic result in rainfall of 3– 4 m p.a. on the more coastal highlands, decreasing steadily inland. The wettest spot in Africa, Cape Debundscha, at the foot of Mt Cameroon, receives 10–15 m p.a., and rainfall there is almost continuous throughout the year. Rainfall in most of the highlands occurs mainly from May to November (Courade 1974, Cable & Cheek 1998). The mountains disrupt the normal bimodal pattern seen in West Africa, resulting in one long, instead of two shorter rainy seasons per annum. Submontane or cloud forest, is generally recognised as occurring in the 800–200 m alt. band and is characterised by the presence of black, humic soils and the presence of abundant epiphytic, pendulous mosses on woody plants (Cheek et al. 2004). Cloud forest once extended from the coastal highlands inland to the Bamenda Highlands at least, but its clearance has been near total in the Bamileke Highlands (now intensively cultivated for *Coffea arabica* L.) and agriculture is also extensive in the Bamenda Highlands where human population is also dense. Total primary forest loss for Cameroon is reported as 47.7% and loss per year from 2001–2018 at 52,272 ha per annum (Mongabay (https://rainforests.mongabay.com/deforestation/archive/Cameroon.htm downloaded 18 July 2019). However, forest loss has not been evenly spread throughout the country, and has been most extensive in high altitude areas above the malaria zone with relatively fertile soils. In the Bamenda Highlands the sole surviving patch of any size is the Bali Ngemba forest reserve (c. 8 km^2^), which is not managed for conservation purposes but for timber extraction. Further south some cloud forest remains in places along the steep west-facing escarpment that links the plateaux with the lowlands, such as at Lebialem, but this is steadily also being cleared for agriculture (Tchiengue in Harvey et al. 2010). The largest surviving blocks of submontane forest in Cameroon are thought to be those in the Bakossi Mts and Mt Kupe where extensive tracts still blanket the mountains, although even her,e clearance upslope for small-holder agriculture continues (Cheek et al. 2004). The cloud forest vegetation and species composition at several of these locations have been characterised in the series of conservation checklists referred to above, and additionally that for Mt Cameroon by Tchouto et al. (1999). However despite this, some areas, such as the Bakossi Mts, remain incompletely sampled, and the Rumpi Hills and Lebialem Highland areas are even more poorly known to science, while some highland areas remain totally unsampled. There is no doubt that many tens of species remain to be discovered and published. Among other cloud forest plant species discovered in cloud forest in the Cameroon highlands in the last 20 years, are (grouped by life-form, in alphabetical order by genus):

#### Herbs

*Brachystephanus oreacanthus* Champl. (Champluvier & Darbyshire 2009), *Coleochloa domensis* Muasya & D.A. Simpson (Muasya et al. 2010), *Costus kupensis* Maas & H. Maas (Maas-van de Kamer et al. 2016), *Dracaena kupensis* Mwachala et al. (Mwachala et al. 2007), *Impatiens etindensis* Cheek & Eb. Fischer (Cheek & Fischer 1999), *Impatiens frithii* Cheek (Cheek & Csiba 2002a), *Isoglossa dispersa* I.Darbysh. (Darbyshire et al. 2011)*, Kupea martinetugei* Cheek & S. A. Williams (Cheek et al. 2003), *Ledermanniella onanae* Cheek (Cheek 2003), *Ledermanniella pollardiana* Cheek & Ameka (Cheek & Ameka 2008), *Mussaenda epiphytica* Cheek (Cheek 2009).

#### Shrubs

*Chassalia laikomensis* Cheek (Cheek & Csiba 2000)*, Coffea montekupensis* Stoff. (Stoffelen et al. 1997), *Memecylon kupeanum* R.D.Stone, Ghogue & Cheek (Stone et al. 2008), *Oxyanthus okuensis* Cheek & Sonké (Cheek & Sonké 2000)*, Psychotria darwiniana* Cheek (Cheek et al. 2009), *Psychotria geophylax* and *P. bakossiensis* Cheek & Sonké (Cheek & Sonké 2005)*, Psychotria kupensis* Cheek (Cheek et al. 2008), *Psychotria moseskemei* Cheek (Cheek & Csiba 2002b).

#### Trees

*Allophylus ujori* Cheek (Cheek & Etuge 2009b)*, Coffea bakossii* Cheek & Bridson (Cheek et al. 2002), *Deinbollia oreophila* Cheek (Cheek & Etuge 2009a)*, Diospyros kupensis* Gosline (Gosline & Cheek 1998), *Dovyalis cameroonensis* Cheek & Ngolan (Cheek & Ngolan 2007), *Microcos magnifica* Cheek (Cheek 2017), *Myrianthus fosi* Cheek (Cheek & Osborne in Harvey et al. 2010), Newtonia *duncanthomasii* Mackinder & Cheek (Mackinder & Cheek 2003)*, Rhaptopetalum geophylax* Cheek & Gosline (Cheek et al. 2002), *Talbotiella bakossiensis* Cheek (Mackinder et al. 2010), *Ternstroemia cameroonensis* Cheek (Cheek et al. 2017) and *Uvariopsis submontana* Kenfack, Gosline & Gereau (Kenfack et al. 2003).

These cases illustrate the scale at which new discoveries are being made in the Highlands of western Cameroon which already contains the most species-diverse degree squares documented in tropical Africa; and which includes several Pleistocene refuge areas (Cheek et al. 2001).

Most of the species listed above are threatened with extinction, since they are narrow endemics with small ranges, restricted to mainly submontane (cloud) forest patches which are steadily being cleared (Onana & Cheek 2011).

The number of species described as new to science each year regularly exceeds 2000, adding to the estimated 369 000 already known (Nic Lughadha et al. 2016), although the number of flowering plant species known to science is disputed (Nic Lughadha et al. 2017). Only 7.2% have been assessed and included on the Red List using the IUCN (2012) standard (Bachman et al. 2019), but this number rises to 21–26% when additional evidence-based assessments are considered, and 30–44% of these assess the species as threatened (Bachman et al. 2018). Newly discovered species, such as that reported in this paper, are likely to be threatened, since widespread species tend to have been already discovered. There are notable exceptions to this rule (e.g. *Vepris occidentalis* Cheek (Cheek et al. 2019) a species widespread in West Africa from Guinea to Ghana). Generally, it is the more localised, rarer species that remain undiscovered. This makes it all the more urgent to find, document and protect such species before they become extinct, as is *Oxygyne triandra* Schltr. (Cheek et al. 2018b), or possibly extinct, in the case of another Cameroon Highland cloud forest tree, *Vepris bali* Cheek (Cheek et al. 2018c). Most of the 815 Cameroonian species Red Listed in the “Red Data Book, Plants of Cameroon” are threatened with extinction due to habitat clearance, mainly for small holder and plantation agriculture following logging (Onana & Cheek 2011). Efforts are now being made to delimit the highest priority areas in Cameroon for plant conservation as Tropical

Important Plant Areas (TIPAs) using the revised IPA criteria set out in Darbyshire et al. (2017). This is intended to help avoid the global extinction of additional endemic species such as *Tricalysia elmar*.

## ACKNOWLEDGEMENTS

We thank Dr Helen Fortune-Hopkins for reviewing an earlier version of this manuscript. The botanical surveys in Cameroon which resulted in this paper were mainly supported by Earthwatch Europe (1993–2005) and by the Darwin Initiative of the UK Government through the Plant Conservation of Western Cameroon, and the Red Data Book of Cameroon projects, both led by RBG, Kew, working with the IRAD-National Herbarium of Cameroon.

Gaston Achoundong, former head of the National Herbarium of Cameroon (YA) and his successors Jean Michel Onana and Florence Ngo Ngwe are thanked for their collaboration and support over the years.

